# Configural properties underlie the perceived *faceness* of a stimulus

**DOI:** 10.1101/509026

**Authors:** Carmine Gnolo, Mario Senden, Alessandro Grillini, Frans W. Cornelissen, Rainer Goebel

## Abstract

The global geometrical arrangement of face parts (eyes over nose over mouth), commonly referred to as first order configural (FOC) properties, is believed to constitute a fundamental aspect of face detection. Indeed, several brain regions in the face processing network have been shown to display increased activity in response to abstract stimuli for face-like as compared to random spatial arrangements. However, the absence of a mathematical formalization of FOC properties makes it difficult to systematically study their contribution to face perception. To fill this gap, we used analytical geometry to define a set of 19 FOC features. In two psychophysical studies, a 2-alternative forced choice task assessing difference in perceived *faceness* between two stimuli and a Likert test assessing measuring perceived *faceness* of individual stimuli, we evaluated the contribution of each feature to the perceived *faceness* of abstract stimuli consisting of four black rectangular shapes reflecting the two eyes, the nose and the mouth. From the two experiments two subsets of features relevant for face detection were selected, including 10 and 11 features, respectively. Interestingly, 7 features are shared between the two sets. The difference between the subsets reflects processing of individual FOC features in the first study and holistic processing in the second study. The partial superimposition, on the other hand, is reflective of a shared basic mechanism in the perception of first order configural properties.

## 1. Introduction

Face processing is one of the most investigated subdomains of high level human vision. One reason for this is the immediate relevance of faces for everyday social interactions and the accompanying evolutionary value of successfully recognizing faces (Pascalis & Kelly, 2009). As such it is hardly surprising that a specialized network in the brain able to parse a high density of information from visual stimuli is dedicated to face processing. Because faces elicit a stronger response than other objects in a number of cortical areas (Duchaine & Yovel, 2015; Kanwisher & Barton, 2011; Liu, Harris, & Kanwisher, 2002; Tsao & Livingstone, 2008), we presume that there are patches of the cortex that engage in face detection by evaluating mid-level visual features and that modulate activation in portions of the ventral stream.

The problem of face detection can be reframed as the identification of those properties of an image that make it face-like. Knowing these properties and being able to modulate them is not only interesting because it may lead to an improved understanding of how face detection works. It may also provide improved means for modulating face-likeness in order to study face-specific processes including their contribution to visual crowding (Sun & Balas, 2014), face preference in newborns (Macchi Cassia, Kuefner, Westerlund, & Nelson, 2006), deviation of attention and gaze in autism (Golarai, Grill-Spector, & Reiss, 2006), and holistic processing (J. W. Tanaka & Sengco, 1997; Tsao & Livingstone, 2008).

Studies exploring face detection tend to use one of two common approaches to face-likeness modulation. The first is visibility modulation (Chen, McBain, & Norton, 2015; Gilad-Gutnick, Yovel, & Sinha, 2012; Jiang et al., 2011), which involves altering low level properties of stimuli (e.g., hue, luminance, contrast) to make faces less recognizable by applying filters, additive noise or phase scrambling. Those approaches neglect the modulation of higher level visual properties of faces (e.g. the disposition of face parts), which lie at the foundation of face detection (Mondloch, Le Grand, & Maurer, 2002). In spite of this, what can make such approaches attractive is that they enable a continuous modulation of faceness. The other approach is categorical modulation (Bayliss & Tipper, 2005; de Haas et al., 2016; Golarai, Ghahremani, Eberhardt, & Gabrieli, 2015; Macchi Cassia et al., 2006), wherein a stimulus is either a realistic face or a face whose components are scrambled which might involve moving them away from their archetypal position, operating a tile scrambling, or degrading the configural properties of the face by simply inverting it. In contrast to visibility modulation, this approach does focus on high level properties of faces, but merely in a categorical manner, thus excluding the possibility for continuous modulation. Relying on these two approaches therefore limits the ability to study the details of the face detection process.

To overcome the shortcomings of existing approaches, a potentially better approach should allow for the continuous modulation of high level properties of faces. Furthermore, it should allow to specifically modulate those properties relevant for face detection. It is well established in the literature that face perception can be subdivided in three main parts: featural processing, involving how each part of the face (eyes, nose, mouth) looks; configural processing, involving the spatial relations among the parts of the face; and holistic processing, integrating all this information together (Maurer, Grand, & Mondloch, 2002; J. W. Tanaka & Gordon, 2011). Configural processing can be further divided into first order and second order relational configural properties. First order configural processing (FOC) concerns the global spatial disposition of face parts (i.e. eyes over nose over mouth) and is relevant for face detection (Sheehan & Nachman, 2014). Second order categorical processing, on the other hand, refers to the detailed metric spacing between face parts such as interocular-distance and is relevant for face recognition (Sheehan & Nachman, 2014). Face detection and face recognition answer to opposite requirements: the former focuses on what is common among faces, while the latter focuses on what makes faces unique.

Several studies on infants and prosopagnosia patients have shown that those two functions are decoupled (Maurer et al., 2002), with FOC processing and face detection still present in people otherwise unable to perform face recognition. A similar distinction between the roles of first and second order configural processing can be obtained when considering pareidolia stimuli. All properties of such stimuli are incoherent with an actual face with the exception of FOC properties, and yet the brain shows similar activation patterns as for real faces (Hadjikhani, Kveraga, Naik, & Ahlfors, 2009). Finally, featural and second level configural properties exhibit extensive phenotypic as well as genomic variability (Sheehan & Nachman, 2014). This is in line with these properties being mainly related to face recognition which benefits from individuals being easily discriminable.

Given these observations, we argue that faceness can be formalized in terms of FOC properties. We developed such a formalization evolving it from a merely categorical description to a computational model based on geometrical properties of a distribution of face parts. Specifically, our formal description is inspired by the reference lines used by artists when drawing faces. That is, we built a system of reference lines which define the relative position and alignment of all face parts to each other. To restrict the information represented in the stimuli to configural only, we built it as a disposition of four black rectangles on a white background. This kind of stimulus enables us to fully represent the configural properties that can be shown among face parts, while excluding all details related to their appearance. The omission of realistic face parts is justified by studies having shown how synthetic stimuli such as cartoon faces, can be used to effectively drive neural responses (Freiwald, Tsao, & Livingstone, 2009; Kobatake & Tanaka, 1994; K. Tanaka, 2003).

Based on this stimulus and the aforementioned concepts of FOC processing, we defined a set of 19 features formalizing the alignment to a geometric archetype. A recent study revealed linear neural tuning to complex geometrical features representing information relevant for face recognition(Chang & Tsao, 2017). This encouraged us to follow a similar modelling approach for face detection. In line with this, we mathematically represent each configuration with a synthetic matrix consisting of only twelve elements. This enables us to easily extract all features using direct formulae instead of relying on complex algorithms. This synthetic representation furthermore enables us to work over a large stimulus space described by a high-dimensional feature space from which configurations with specific properties can be directly selected for behavioural experiments.

Here, we conducted two experiments, a 2-alternative forced choice experiment as well as a Likert rating experiment, to evaluate our model in terms of individual face features as well as their holistic integration. We find that two sets of respectively 10 and 11 features (with 7 features in common) can sufficiently account for perceived faceness.

## 2. General method

### 2.1 Stimulus space

Prior to defining the features related to FOC properties, it is necessary to construct a ***stimulus space*** that allows for the efficient description of configural pictures, the extraction of features and the generation of stimulus sets with specific configural properties. The first step of this procedure relates to the choice of how to represent face parts. We limit our descriptions to face stimuli consisting of eyes, nose and mouth. Using a gradual simplification of the stimulus, similar to the approach taken in (Kobatake & Tanaka, 1994), we started from realistic drawings, which would still contain information on the detailed look of each part (commonly referred to as *featural* information) and stripped off information until we were left with four black rectangles of equal size on a white background. The only information represented in these stimuli is the position and alignment of each part. Given this, it is possible to fully represent configural information in terms of position and alignment while excluding any detail regarding each individual part. This choice is especially convenient because it allows for changing the face parts while retaining the configural description, using the rectangles as bounding boxes for more detailed face parts.

To keep the representation as general as possible, the only assumptions are 1) that the whole face is enclosed within a square and 2) that all face parts are enclosed within rectangles of equal size. Given this, we can define a frame of reference in Cartesian coordinates. The face parts are located inside the subspace (*x, y*) ∈ [0,1] × [0,1]. In this way it is easy to describe any squared picture, normalizing all distances to the size of its side. We associate each of the parts with a unit vector, applied to the centre of the part and aligned to the x axis in an archetypal configuration. Starting from this, we can fully describe the geometrical properties of each part knowing the origin of the unit vector and its angle with respect to the *x* axis. The angles are assumed positive in the counter clockwise direction. In our specific case, where face parts are black rectangles, we restrict the value of the angle to 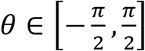 because of rotational symmetry.

Using this formalization, each configuration can be fully described using three values for each part; i.e. its coordinates and angle, leading to the 4 × 3 matrix:

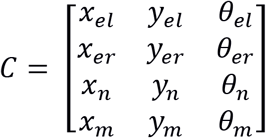

where *el* indicates the left eye, *er* the right eye, *n* the nose and *m* the mouth. We refer to this as the *configural matrix of a stimulus*. This representation of the configurations is especially convenient compared to full-fledged images because, in contrast with the at least thousands of pixels (e.g. 150 x 150) that would be needed to represent a whole image, it fully describes all information used by the model with 12 numbers and is by definition invariant to the appearance of the parts.

### 2.2 Configural model

Configural properties are bound to the position and alignment of the individual parts constituting a face. This kind of information can be naturally described using geometry. Therefore, a first step to go beyond the categorical description of FOC properties of faces is to identify and formalize a set of geometrical relationships among those parts common to all faces. Artists often use geometrical constructions as an aid to draw complex objects, including faces (see figure 1). Within such constructions, the parts of a face are aligned with a cross defined by the line passing through the centre of the eyes and a perpendicular line passing through the mid-point between the eyes. We base our model of FOC on this concept, defining three reference axes and using measures of misalignment of face parts to those axes as features. The amount of misalignment of a single rectangle determines how well it fits in the whole face structure, making it more difficult to recognize it as a specific face part. The misalignment between axes measures the destabilization of the whole structure of the face and reflects destruction of the symmetry of the positions.

**Figure 1:**
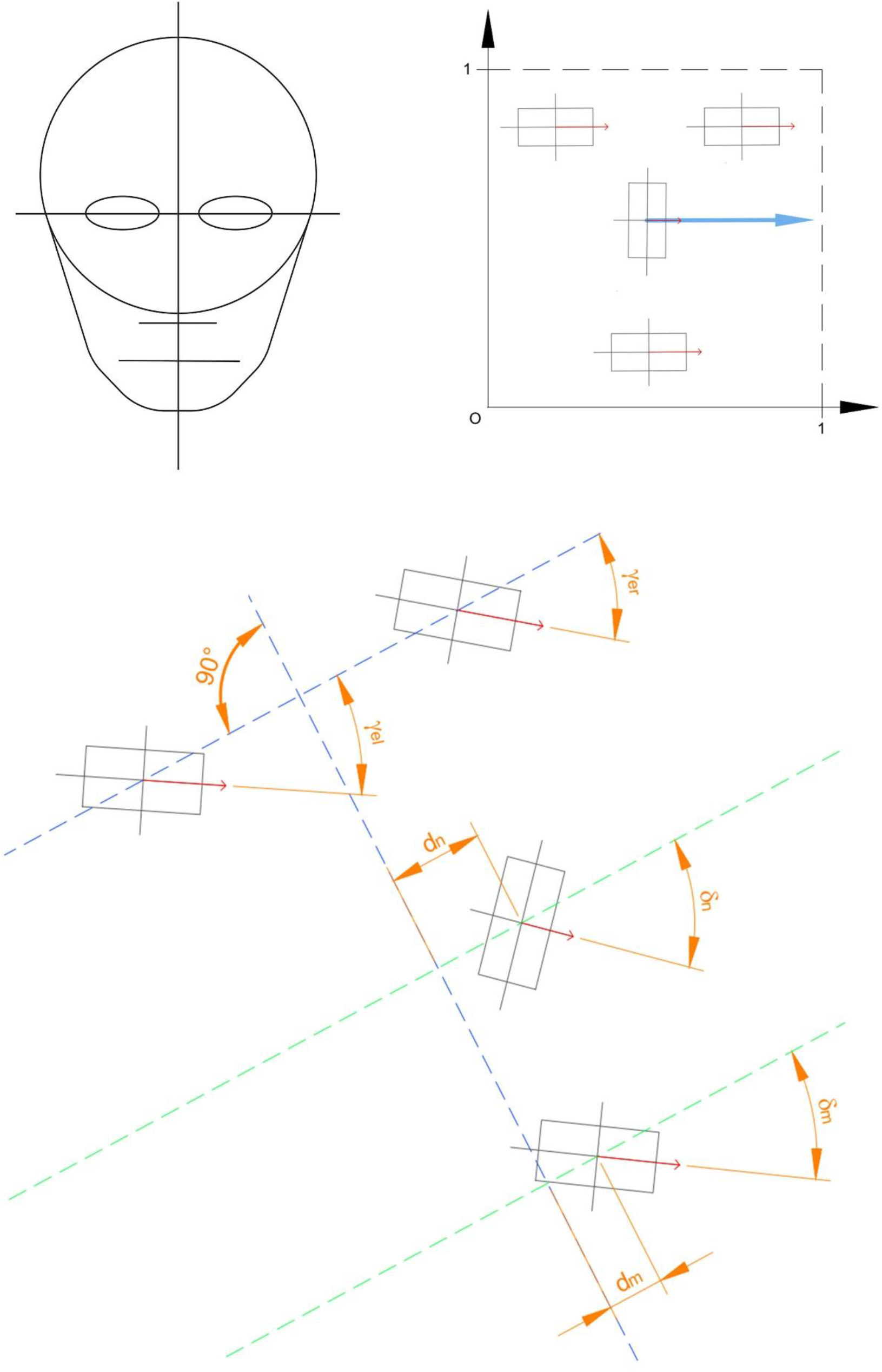
Definition of face axes. **(A)** Schematic outline used by artists to draw faces. **(B)** Archetypal face configuration located in the representation space defined by the artists’ outline. **(C)** Schematic overview of the features that can be defined based on this formalization (in orange).

The geometric construction is based on three main lines, determined by the position of some of the parts. Because all shapes are identical, it is only their position and orientation and their relation to one another that makes them recognizable as a specific part (e.g. the one in the centre is usually the nose). This implies that parts which deviate too much from their typical location might be reinterpreted as different face parts. To prevent this, we limit the range in which the parts are displaced from an archetypical disposition. Once the geometrical model is conceptually established, we formalize it mathematically to explicitly extract features using direct formulae applied to the elements of the configural matrix. Reference lines are described with line equations and parts are represented by vectors given by the coordinates of their application point and their alignment. We use concepts from analytical geometry and trigonometry to measure distances and alignments using those line equations and vectors. We express all equations of our lines in the explicit form

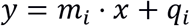

where *i* indicates the line to which the slope *m_i_* and the intercept *q_i_* belong. This way all lines are uniquely identified by the couple (*m_i_*, *q_i_*). We begin the geometrical construct by drawing a line passing through the centres of the eyes:

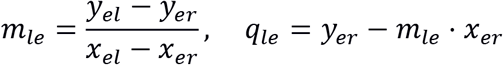

where *l_e_* indicates line of the eyes.

After this we define the point *P* as the midpoint between the eyes

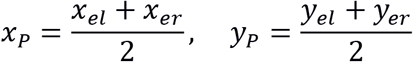

Given *P*, we define our first main axis, *l_pe_*, as perpendicular to *l_e_* and passing through the point *P*:

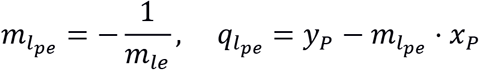

The second main axis is the line *l_y_*, parallel to the *y* axis and passing through P, with equation

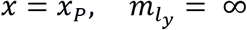

The third main axis *l_nm_* is the line passing through the centres of the nose and mouth:

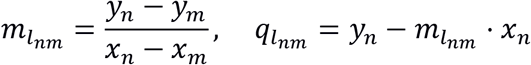

The features in the model will measure misalignment and distance of the face parts from three main axes.

**Figure 2:**
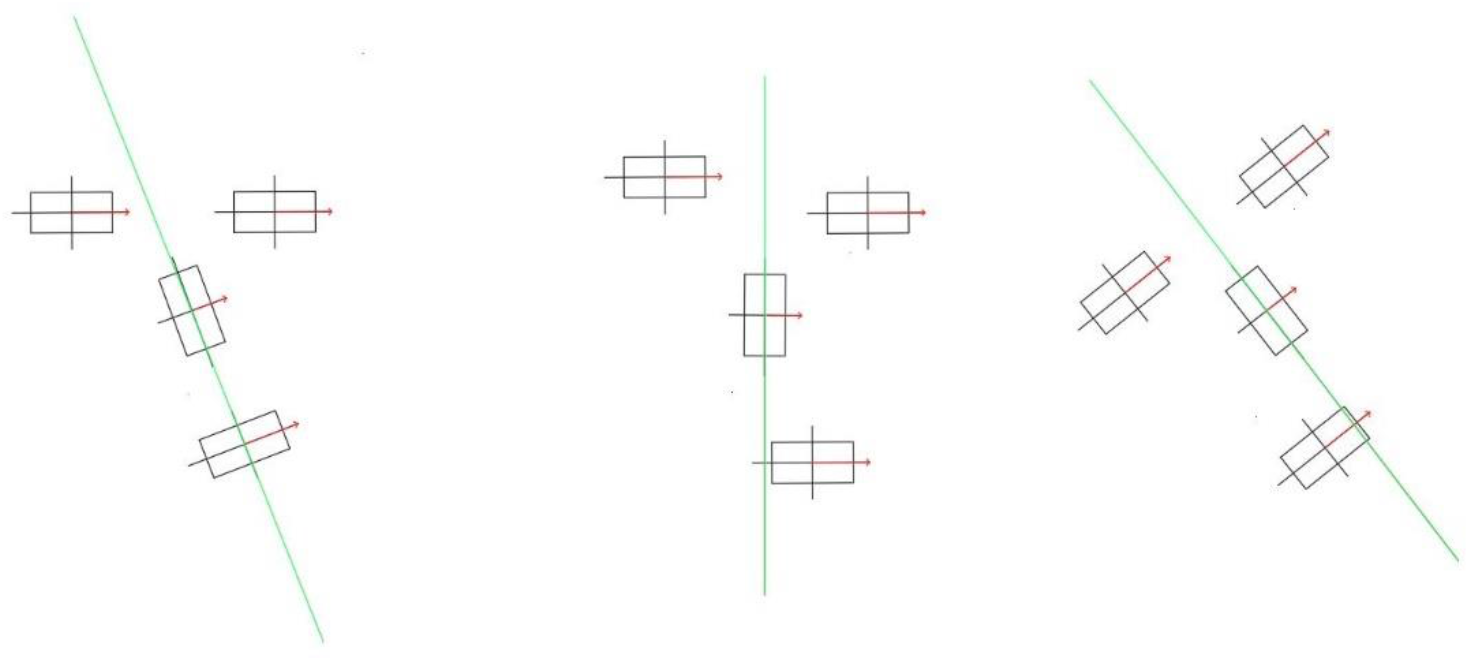
Examples of three different dominant directions **(A)** A face whose dominant direction is defined by the line passing through the centres of the nose and the mouth (*l*_nm_). **(B)** A face whose dominant direction is parallel to the y-axis (the *l_y_*). **(C)** A face whose dominant direction is given by the line perpendicular to the alignment of the eyes (*l_pe_*).

To measure misalignments, we define some quantities we refer to as *angular deviations*, *adev*(*θ*_1_, *θ*_2_), where *θ* is the angle between the vector representing either a face part or a line and the positive direction of the *x* axis.

The parts we use in this case are rotationally symmetric by π radians; thus, *adev* needs to show this periodicity:

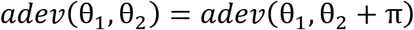

Furthermore, we are interested in the magnitude of the misalignment but not its direction. This means that the maximum angular deviation occurs when parts are perpendicular. Finally, we restrict the values of all features to the range from 0 (minimal deviation) to 1 (maximal deviation):

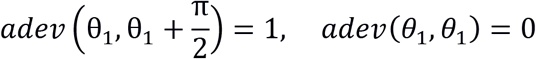

with

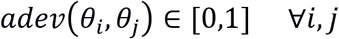

and ensure that the sign does not change with the direction of angular deviation:

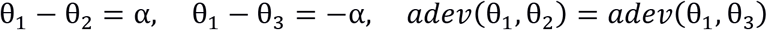

To achieve this, it is convenient to rewrite the angles as the phase of complex numbers. In this representation differences between angles can be expressed as movements on the goniometric circle, which are periodical. We can adapt this periodicity to meet our needs, which in our case is a periodicity of *π*/2. After that we normalize to have values in [0,1]:

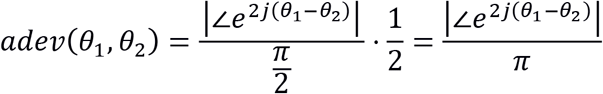

This also works if one of the angles *θ* is expressed as the *atan* of an angular coefficient *m*, we just assumed the two interchangeable for clarity. The simplest case for angular deviation between parts and lines is with respect to *l_y_*: in this case the deviation is just the normalized modulo of the alignment of the parts with the x axis

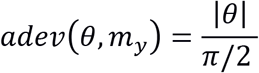

To compute the distance of a point *P*_0_ = (*x*_0_,*y*_0_) from *l_pe_* we used the formula:

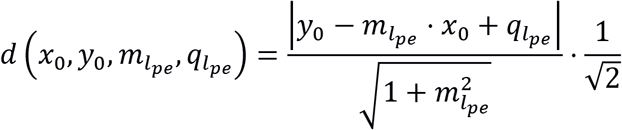

In case the line was parallel to the y axis, this would have resulted in an indeterminate form, because *m_l_* = ∞. In those cases, we used another formula which exploits the fact that *l_pe_* passed through the point *P*:

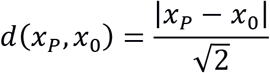

Those distances were divided by 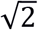 because that is the largest distance in the 2-dimensional domain in which the parts are represented, corresponding to the diagonal of the square. In the case of *l_y_*, this axis was parallel to the *y* axis by definition and passing through *P*. We could then always apply the second formula.

The last two features were more complex and were computed using the results from the other features. They relied on the concept of *dominant direction:* we assumed that when perceiving a face in a configuration we would always perceive it oriented along one of the main axes that we described, *l_y_ l_pe_* or *l_nm_*, and that this dominant axis would be the one exhibiting the least deviation from all the other features related to it. Note that we refer to direction and not alignment because this will be defined by the vertical disposition of the parts. The main problem with this approach is that by definition it is not possible to compute the distance of the mouth and the nose from *l_nm_* because it would always be zero. Moreover, distances contribute more to the decline of faceness than alignments. We account for this by multiplying the alignment deviation values by a coefficient *k* = 1.2, which was determined by visually evaluating the dominant direction perceived with the one computed using different values for a set of test configurations. While the method to define this coefficient may be somewhat subjective, it gave us the possibility to define a dominant direction with reasonable precision and compute vertical disruption and alignment without influencing the other features.

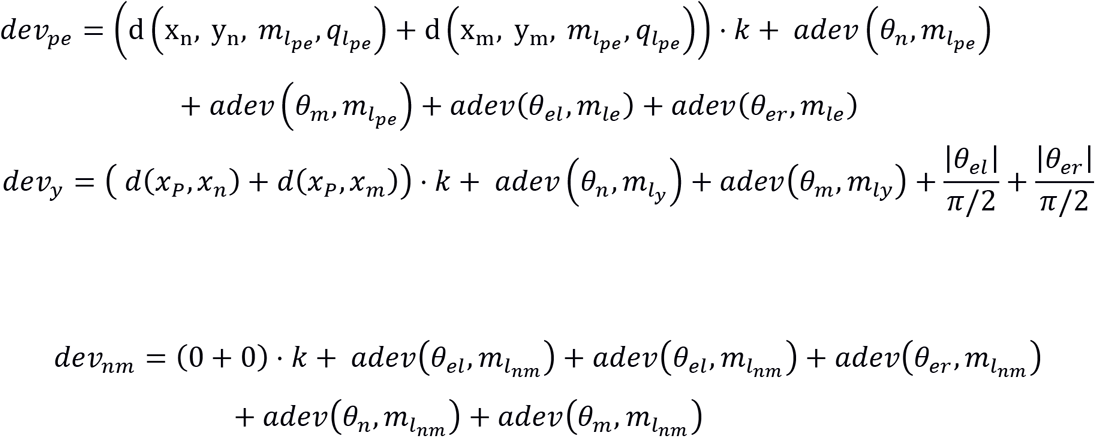

Once we determined which of these deviation scores was lowest, we regarded the corresponding axis as dominant. The direction used to describe the alignment of the configuration was defined as the one perpendicular to the dominant axis. Similar to what we did for individual face parts, we defined the orientation of the whole configuration as a unit vector whose angle *θ_D_* is 0 radians when the face is oriented according to the archetypal configuration. In this case there is no *π* rotational symmetry and the orientation can be defined in the range [0,2*π*]. To evaluate the face orientation *θ_f_*, we rotate the configuration until *l_e_* is parallel to the *x* axis, rotating the configuration by an angle − *atan*(*m_le_*) (recall that this has codomain in 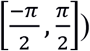 because the vertical disposition of the parts always depends on *l_pe_*, independently from the dominant axis: *l_e_* divides the plane in two areas, one over the eyes and the other under them, so the vertical order must be evaluated along its perpendicular *l_pe_*. There are two possible cases that correspond to a valid face: in a straight face, the order of the parts is eyes – nose – mouth, on an inverted one it is mouth – nose – eyes. Following this principle, in the first case *θ_f_* = *θ_D_*, in the second case it would be *θ_f_* = *θ_D_* + *π*.

We defined a binary feature called vertical disruption, to which we assign a value of 0 if the order of parts corresponds to one of the previously described cases, and a value of 1 otherwise. This represents an immediate extension of the former concept of categorical first order configural properties, applied to our new framework. In those cases, we assume *θ_f_* = *θ^D^*.

The last feature we define is inversion, which we compute from the orientation of the configuration:

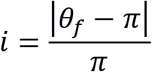

This way the feature will have maximum value of 1 for an orientation of *π* radians, corresponding to an inverted face, and minimum value of 0 for an angle of 0 radians, with the magnitude independent from the direction of the deviation.

**Table 1:** Features. We refer to the features using the ID number in the first column. The last column indicates in which subset the feature was selected: 2-AFC stands stepwise selection based on the 2-alternative forced choice task, L for the subset determined using the Likert test, and the hyphen indicates that the feature was not selected by any model selection method.

| # | Feature | Description | Subset |
| --- | --- | --- | --- |
| 1 | $adev(\theta_{el}, m_{le})$ | Angular deviation between $l_e$ and the left eye | L |
| 2 | $adev(\theta_{er}, m_{le})$ | Angular deviation between $l_e$ and the right eye | L, 2-AFC |
| 3 | $d(x_n, y_n, m_{l_{pe}}, q_{l_{pe}})$ | Distance between the nose and $l_{pe}$ | L, 2-AFC |
| 4 | $d(x_m, y_m, m_{l_{pe}}, q_{l_{pe}})$ | Distance between the mouth and $l_{pe}$ | 2-AFC |
| 5 | $adev(\theta_n, m_{l_{pe}})$ | Angular deviation between $l_{pe}$ and the nose | 2-AFC |
| 6 | $adev(\theta_m, m_{l_{pe}})$ | Angular deviation between $l_{pe}$ and the mouth | L |
| 7 | $adev(\theta_n, m_{l_y})$ | Angular deviation between $l_y$ and the nose | - |
| 8 | $adev(\theta_m, m_{l_y})$ | Angular deviation between $l_y$ and the mouth | L, 2-AFC |
| 9 | $d(x_p, x_n)$ | Distance between the nose and $l_y$ | L, 2-AFC |
| 10 | $d(x_p, x_m)$ | Distance between the mouth and $l_y$ | L, 2-AFC |
| 11 | $adev(\theta_{el}, m_{l_{nm}})$ | Angular deviation between $l_{nm}$ and the left eye | - |
| 12 | $adev(\theta_{er}, m_{l_{nm}})$ | Angular deviation between $l_{nm}$ and the right eye | - |
| 13 | $adev(\theta_n, m_{l_{nm}})$ | Angular deviation between $l_{nm}$ and the nose | L |
| 14 | $adev(\theta_m, m_{l_{nm}})$ | Angular deviation between $l_{nm}$ and the mouth | - |
| 15 | $adev(m_{l_{pe}}, m_{l_{nm}})$ | Angular deviation between $l_{pe}$ and $l_{nm}$ | L, 2-AFC |
| 16 | $adev(m_{l_y}, m_{l_{nm}})$ | Angular deviation between $l_y$ and $l_{nm}$ | 2-AFC |
| 17 | $adev(m_{l_y}, m_{l_{pe}})$ | Angular deviation between $l_y$ and $l_{pe}$ | 2-AFC |
| 18 | Vertical disruption |  | L, 2-AFC |
| 19 | Inversion |  | - |

## 3. Experiment 1: Sensitivity to the variation of a single feature

### 3.1 Participants

The cohort consisted of 13 healthy participants (average age 25 years, sd 3.6, 3 males). All participants had normal or corrected-to-normal visual acuity. The ethics board of the Faculty of Psychology and Neuroscience at Maastricht University approved the study protocol. All participants provided written informed consent prior to participation.

### 3.2 Stimuli and procedure

To investigate the contribution of each feature towards the perception of the whole configuration as a face, we designed a 2-alternative forced choice (2-AFC) psychophysical experiment. Participants were instructed to fixate a black cross at the centre of a white screen during the resting condition. During choice trials, two face configurations were presented simultaneously at a distance of 5 degrees of visual angle to the left and right of the fixation cross. After freely looking at those, participants needed to indicate which of the two configurations looked most like a face using the left and right arrow keys. They return to fixation during the rest condition. Stimulus presentation and inter-stimulus interval were both set to 1 s. Stimulus presentation was interrupted earlier in case of a response faster than 1 s. This procedure was repeated for 200 trials for each run, with one run per feature, for a total of (200 x 19 = 3800 trials). All participants had a break of 10 minutes after the first 9 runs. The order of runs was counterbalanced in a pseudorandom fashion across participants.

To create a set of configurations per run, we first generated a sample of ~10^9^ configural matrices establishing ranges of positions in which each part could be moved, starting from an archetype. All configurations for which parts overlapped were removed. We then computed the feature values for all matrices as well as the normalized sum of all features. Within each run the feature of interest (FOI) varied while the total faceness due to the remaining features was held constant. That is, we selected only those configurations from the set which showed a normalized sum of the remaining features within the range 0.2 ± 0.03. For the resulting set of configurations, we then identified those two configurations for which the FOI was maximal and minimal, respectively, and generated a linearly spaced vector which spanned this interval in six steps. For each of the six levels we sampled 36 matrices and generated configurations with a value of the FOI as close as possible to the desired level. We then chose one of those as the configuration to use for the level, taking care not to consider images that would cause ambiguity in the interpretation of the parts, judged based on direct observation of the configuration. This was done for all levels and all features.

The criterion used for the configurations sampling to run the psychophysics is shown in Figure 3. The feature of interest is varied covering the widest range possible while keeping the other features linearly independent and their normalized sum as constant as possible.

**Figure 3:**
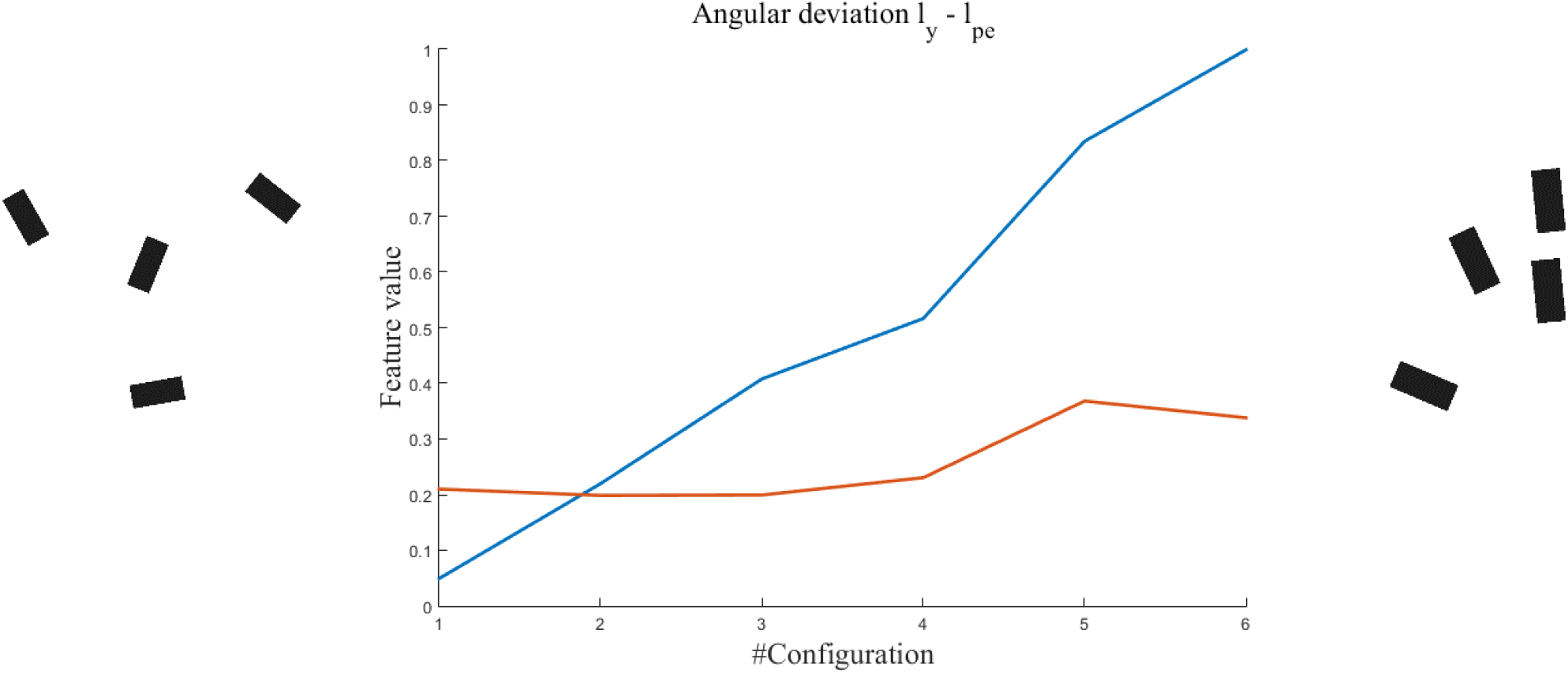
Values of the feature of interest (in blue) and of the normalized sum of the remaining features (in red), for the six configurations used for the psychophysics for one of the features. On the sides are configurations reflective of the minimum (left) and maximum (right) values of the feature.

### 3.3 Materials

The experiment was conducted using MATLAB (version R2015b; The Mathworks Inc.; Natick, MA, USA) and the stimulation suite Psychtoolbox-3 (Brainard, 1997; Kleiner et al., 2007; Pelli, 1997). We displayed the stimuli on a Neovo F-419 monitor, 19 inches, with a refresh rate of 75 Hz. We conducted the experiment in a silent room with low light. The distance of the subject from the monitor was 60 cm during the task, controlled by positioning the subject on previously measured reference lines.

### 3.4 Statistical analysis

The data was analysed using the MATLAB (version R2015b; The Mathworks Inc.; Natick, MA, USA) software tools. Psychometric curves were fitted using logistic regression for both single subject and group data. The group curves were computed by concatenating the subject-specific data.

### 3.5 Results

Figure 4 shows the psychometric curves measured for all the features. Panel (a) shows the fits for each subject while Panel (b) shows the results of the fixed effect group analysis. Some of the features show a very clear sigmoidal pattern (features 9, 4 and 17), whereas for others it is evident that the logistic regression failed (features 1, 6, 7). We show in Table 2 the statistical results of the psychometric curves. We can see that feature 9, angular deviation between *l_y_* and the nose, is not only fit well by the logit, but also shows the strongest perceived effect (estimated via *β*_1_). The features 4, 5, 17, 10 and 15, which come in order of effect strength after feature 9, also show a sigmoid shape. The psychometric curves for the experiments show a goodness of fit and a sigmoid shape comparable to what is usually observed in experiments involving lower level visual features, whose perceptive mechanisms are better known. Similarly, well-fitting psychometric curves are rarely observed for higher level features like the ones presented here and especially in experiments with a design as simple as the one we used. This means that the changes in the features we are proposing here are perceived very clearly by the subject when instructed to only look for faceness. This strongly suggests that those features are relevant to the perception of faceness.

**Table 2:** Statistics related to the psychometric curves

| # | Feature | $\beta_1$ | $SE_{\beta_1}$ | $\chi^2_{2798}$ | $p$ |
| --- | --- | --- | --- | --- | --- |
| 9 | Distance between the nose and $l_y$ | 7.71 | 0.84 | 1324.74 | $\ll 0.001$ |
| 4 | Distance between the mouth and $l_{pe}$ | 3.85 | 0.59 | 462.05 | $\ll 0.001$ |
| 5 | Angular deviation between $l_{pe}$ and the nose | 2.84 | 0.47 | 451.46 | $\ll 0.001$ |
| 17 | Angular deviation between $l_y$ and $l_{pe}$ | 2.58 | 0.19 | 756.04 | $\ll 0.001$ |
| 10 | Distance between the mouth and $l_y$ | 2.37 | 0.57 | 156.47 | $\ll 0.001$ |
| 15 | Angular deviation between $l_{pe}$ and $l_{nm}$ | 1.98 | 0.28 | 499.08 | $\ll 0.001$ |
| 18 | Vertical disruption | 1.86 | 0.22 | 824.19 | $\ll 0.001$ |
| 16 | Angular deviation between $l_y$ and $l_{nm}$ | 1.50 | 0.48 | 192.42 | $\ll 0.001$ |
| 3 | Distance between the nose and $l_{pe}$ | 1.38 | 0.65 | 149.15 | $\ll 0.001$ |
| 8 | Angular deviation between $l_y$ and the mouth | 1.08 | 0.54 | 83.13 | $\ll 0.001$ |
| 2 | Angular deviation between $l_e$ and the right eye | 0.85 | 0.20 | 94.67 | $\ll 0.001$ |
| 11 | Angular deviation between $l_{nm}$ and the left eye | 0.73 | 0.15 | 60.17 | $\ll 0.001$ |
| 14 | Angular deviation between $l_{nm}$ and the mouth | 0.59 | 0.29 | 59.35 | $\ll 0.001$ |
| 12 | Angular deviation between $l_{nm}$ and the right eye | 0.17 | 0.22 | 5.46 | 0.020 |
| 7 | Angular deviation between $l_y$ and the nose | -0.15 | 0.26 | 2.23 | 0.136 |
| 19 | Inversion | -0.75 | 0.41 | 80.30 | $\ll 0.001$ |
| 13 | Angular deviation between $l_{nm}$ and the nose | -0.75 | 0.25 | 66.49 | $\ll 0.001$ |
| 1 | Angular deviation between $l_e$ and the left eye | -0.89 | 0.31 | 111.41 | $\ll 0.001$ |
| 6 | Angular deviation between $l_{pe}$ and the mouth | -1.38 | 0.24 | 145.55 | $\ll 0.001$ |

Note. Features are ordered by decreasing $\beta_1$ . Furthermore, $\beta_1$ stems from group data while the standard error of measurement of $\beta_1$ was computed from individual subject analyses.

**Figure 4:**
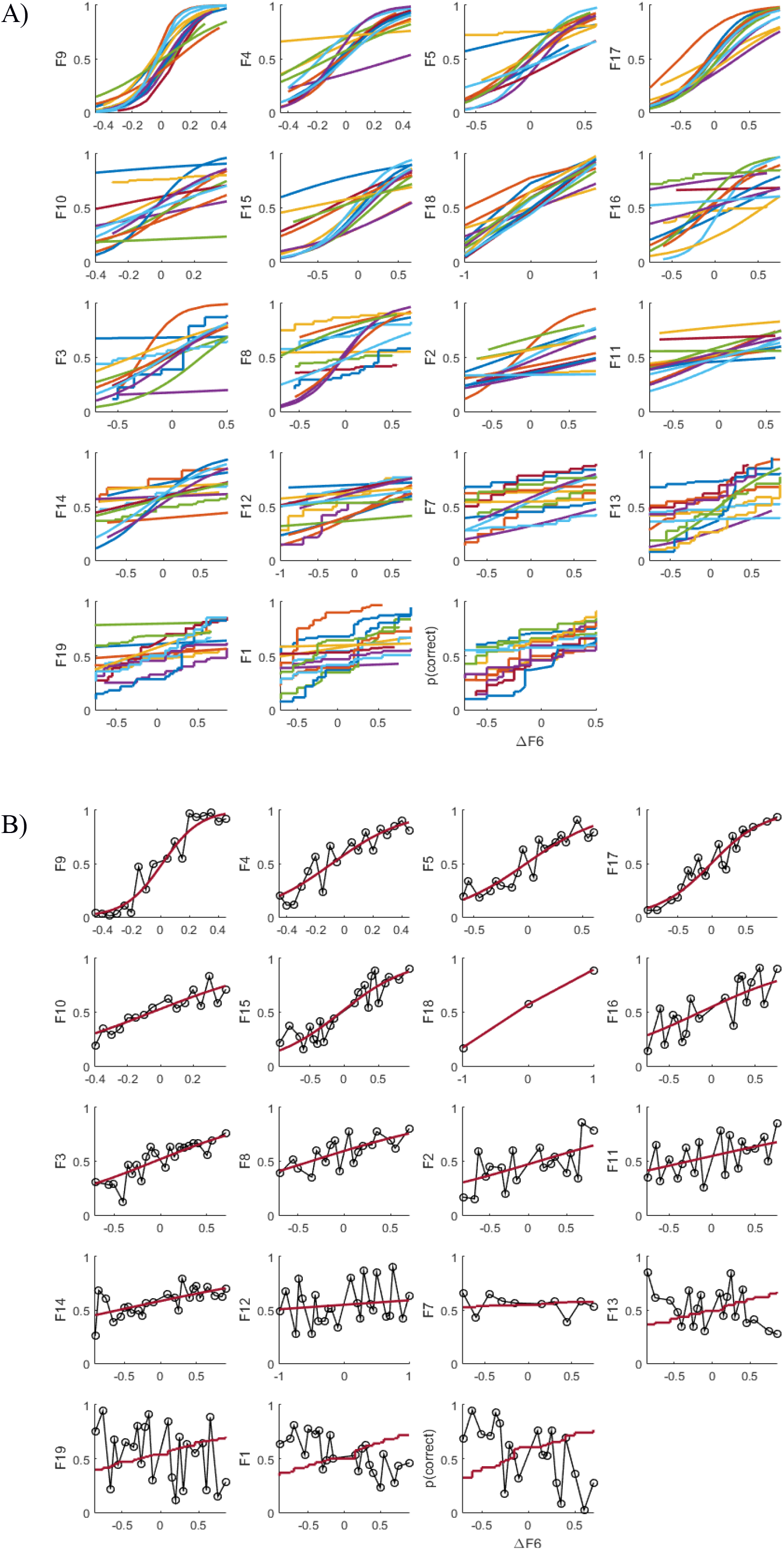
Psychometric curves obtained from the 2-AFC experiment. **(A)** Fitted curves for each subject per feature (subjects differ in color). **(B)** Group-level curves per feature. Fitted curves, shown in red, are based on the data from all the subjects put together (shown in dotted black). The features are ordered by decreasing *β*_2_ value. The x axes represent the difference between the values of the feature of interest between the left and the right configuration. The y axes represent the probability of correct answer for the specific feature considered. We indicate for each graph which feature is represented using the id numbers shown in Table 1.

## 4. Experiment 2: Evaluation of variation of multiple features

### 4.1 Participants

The cohort consisted of 29 healthy subjects (average age 33 years, sd 8.1, 7 females), recruited online. The ethics board of the Faculty of Psychology and Neuroscience at Maastricht University approved the study protocol.

### 4.2 Stimuli and procedure

The experiment consisted of a simple Likert scale task in which participants were shown a configuration and asked to rate how much it resembled a face on a scale from 0 (not *facelike* at all) to 9 (completely *facelike*). This task was repeated for 259 configurations, chosen to have a multivariate Gaussian distribution of feature values, following a sampling procedure described in Appendix A.

### 4.3 Materials

The online surveying platform Qualtrics (Qualtrics, Provo, UT) was used to administer the test to participants, who viewed the stimuli on their own devices.

### 4.4 Statistical analysis

We used MATLAB (version R2015b; The Mathworks Inc.; Natick, MA, USA) to analyse the data from Qualtrics. We performed model selection based on the Bayesian Information Criterion, with the logarithm of the faceness scores as dependent variable and the sum of the features as predictor. We used the logarithm of the faceness rating because we expected it to follow the Weber-Fechner law and hence be proportional to the logarithm of the stimulus. We kept the model linear by ignoring the contribution of the interactions between features. Note that because the features measure deviation from the archetype (degradation of the faceness) while the model measures faceness, regression coefficients are negative. The algorithm reduced the cardinality of the features space, selecting a subset of the most relevant ones.

To select a subset of features to which subjects would be maximally sensitive while simultaneously assuring that the subset is able to adequately describe global faceness, we decided to additionally perform a custom stepwise regression using data from both experiments. That is, we ranked the features based on the strength of their effect, measured as the *β*_1_ value, in the psychometric experiment. Then we fitted the Likert scale data using a model including progressively more features following this ranking. We stopped adding features once the model fit was sufficiently large (*R*^2^ ≥ 0.8).

### 4.5 Results

In Figure 5 we show a 2D representation of the fit between the logarithm of the Likert scale data and the model built using the features selected by the Bayesian Information Criterion. The subset of most predictive features consists of 10 elements and can explain most of the variance in the data (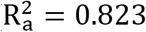, *F*(_11,248_) = 115, *p* ≪ 0.001). The three most predictive features are the distance between the nose and l_pe_ (*t*_11_ = −4.01, *p* ≪ 0.001), the distance between the mouth and l_y_ (*t*_11_ = −6.35, *p* = 0.00024) and the angular deviation between the *l_y_* and the nose (*t*_11_ = −2.39, *p* = 0.018). We show the details about those fittings in Table 3.

**Figure 5:**
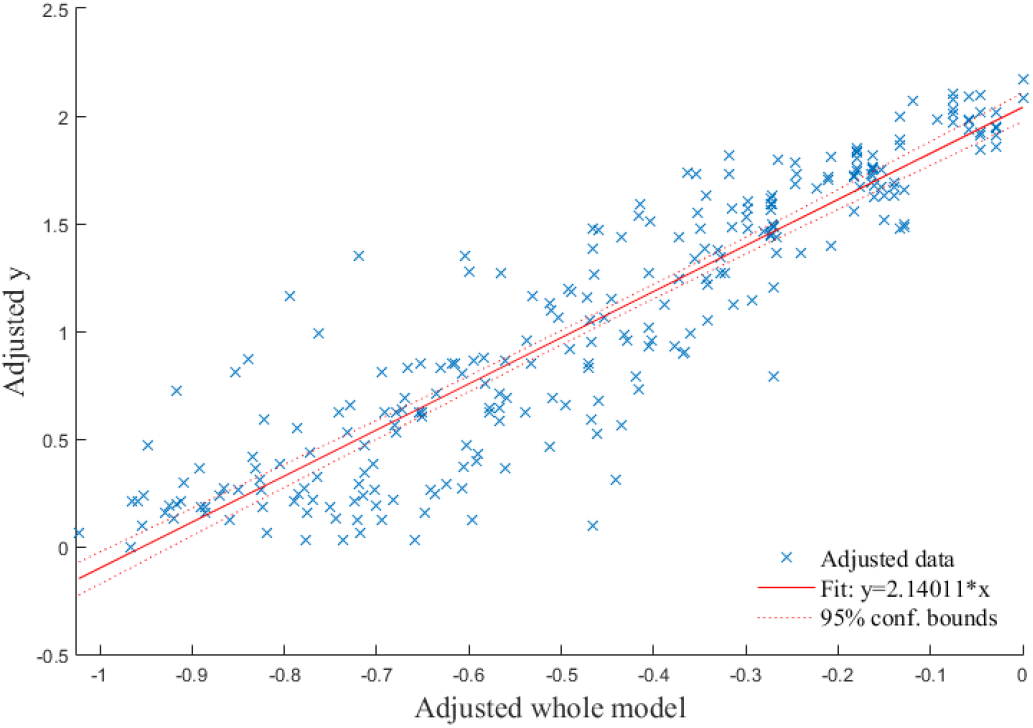
2D representation of the multidimensional regression based on the Likert data, realized using the Frisch-Waugh-Lovell theorem

Figure 6 shows how *R*^2^ varies as features are added to the model following the ranking based on *β*_1_ values from experiment 1: it approximately plateaus after the 11^th^ feature, when it reaches a *R*^2^ value of 0.812. The subsets selected by the Likert analysis alone and the ones which combine 2-AFC and Likert share 7 features.

**Figure 6:**
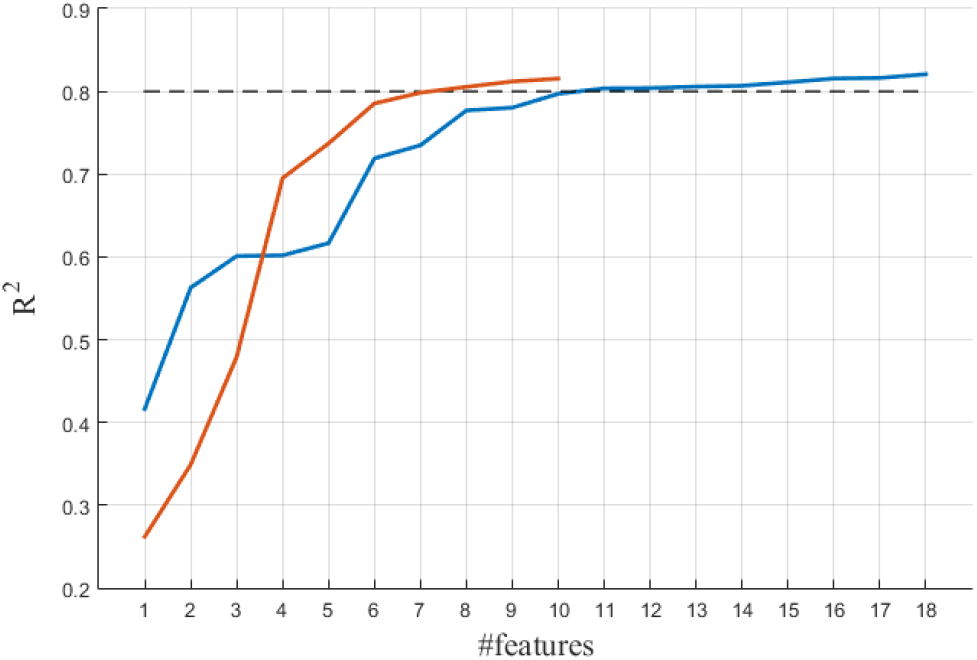
Values of *R*^2^ depending on the number of features included in the subset used for the regression. In orange are the results for the stepwise regression based on the Bayesian Information Criterion. In blue are the values obtained using the *β*_1_ based stepwise regression. The red line stops at 10 features because the stepwise algorithm has reached its stopping criterion.

**Table 3:** Fitting parameters estimates and statistics for the model obtained from *β*_1_ based model selection

| # | Feature name | Estimate | SE | $t_{11}$ | $p$ |
| --- | --- | --- | --- | --- | --- |
| - | Intercept | 2.04 | 0.04 | 48.09 | $\ll 0.001$ |
| 3 | Distance between the nose and $l_{pe}$ | -1.29 | 0.32 | -4.01 | $\ll 0.001$ |
| 10 | Distance between the mouth and $l_y$ | -1.21 | 0.19 | -6.35 | $\ll 0.001$ |
| 11 | Distance between the nose and $l_y$ | -0.73 | 0.30 | -2.39 | 0.017 |
| 15 | Angular deviation between $l_{pe}$ and $l_{nm}$ | -0.58 | 0.08 | -7.68 | $\ll 0.001$ |
| 13 | Angular deviation between $l_{nm}$ and the nose | -0.51 | 0.07 | -6.80 | $\ll 0.001$ |
| 6 | Angular deviation between $l_{pe}$ and the mouth | -0.34 | 0.09 | -4.04 | $\ll 0.001$ |
| 18 | Vertical disruption | -0.30 | 0.06 | -5.21 | $\ll 0.001$ |
| 8 | Angular deviation between $l_y$ and the mouth | -0.23 | 0.09 | -2.72 | 0.007 |
| 2 | Angular deviation between $l_e$ and the right eye | -0.20 | 0.06 | -3.21 | 0.002 |
| 1 | Angular deviation between $l_e$ and the left eye | -0.12 | 0.05 | -2.38 | 0.018 |

**Table 4:** Fitting parameters estimates and statistics for the model obtained from based model selection

| # | Feature name | Estimate | SE | $t_{12}$ | $p$ |
| --- | --- | --- | --- | --- | --- |
| - | Intercept | 1.95 | 0.04 | 49.60 | $\ll 0.001$ |
| 3 | Distance between the nose and $l_{pe}$ | -0.94 | 0.36 | -2.64 | $\ll 0.001$ |
| 10 | Distance between the mouth and $l_y$ | -0.86 | 0.23 | -3.69 | $\ll 0.001$ |
| 9 | Distance between the nose and $l_y$ | -0.79 | 0.34 | -2.33 | 0.021 |
| 16 | Angular deviation between $l_y$ and $l_{nm}$ | -0.63 | 0.10 | -6.00 | $\ll 0.001$ |
| 15 | Angular deviation between $l_{pe}$ and $l_{nm}$ | -0.52 | 0.10 | -5.31 | $\ll 0.001$ |
| 8 | Angular deviation between $l_y$ and the mouth | -0.35 | 0.09 | -4.11 | $\ll 0.001$ |
| 17 | Angular deviation between $l_y$ and $l_{pe}$ | -0.26 | 0.14 | -1.90 | $\ll 0.001$ |
| 5 | Angular deviation between $l_{pe}$ and the nose | -0.23 | 0.10 | -2.28 | 0.023 |
| 18 | Vertical disruption | -0.22 | 0.07 | -3.26 | $\ll 0.001$ |
| 2 | Angular deviation between $l_e$ and the right eye | -0.19 | 0.06 | -3.04 | 0.003 |
| 4 | Distance between the mouth and $l_{pe}$ | -0.09 | 0.25 | -0.37 | $\ll 0.001$ |

## 5. Discussion

The present paper presents a mathematical formalization and detailed description of first order configural properties related to faceness. Using a geometrical construction inspired by the reference lines used by artists it is possible to define 19 continuous features. Importantly, by evaluating these features experimentally using two psychophysical studies using a 2-AFC paradigm and a Likert test to judge faceness of single instances, we were able to identify those features relevant for the perception of faceness in abstract stimuli. Specifically, our results revealed that 14 features contribute to the perception of faceness. Interestingly, the two experiments had only seven features in common while four features were uniquely relevant in the 2-AFC experiment and three in the Likert test. This is likely because the two experiments highlight different cognitive aspects. The 2-AFC test measures the sensitivity to the variation of individual features when the stimulus is generally not particularly face-like. The Likert test, on the other hand, shows the contribution of individual features when a stimulus can be globally perceived as face-like. Furthermore, the Likert experiment tends to highlight angular deviations of single parts with respect to the line defined by these parts. For instance, it highlights the deviation of each eye from the line between the eyes. Please note that the model is based on relationships between all face parts. This implies that even if some features show more pronounced effects in isolation, they nevertheless have an impact on holistic faceness perception.

Our study provides a parametric model of faces defined by the weighted sum of two sets of either 10 or 11 high-level configural visual features: one is focused on the perception of the variation of a single feature and the other is bound to the global faceness. Those two subsets can be used for different kinds of experiments: the global faceness model can be used to study phenomena related to the faceness of a stimulus as a whole, while the set based on the sensitivity to a single feature can be used to study the neural correlates of each feature.

In the set of 7 features selected by both experiments, we can see the relevance of angular deviations between the major axes (e.g. the angle between l_pe_ and l_mn_) and the measures of distance of the face parts from these axes (e.g. distance of the nose from l_pe_). Those measures have an especially strong effect because they capture the compounding effect of the displacement of multiple parts at once towards the degradation of faceness; that is, they effectively destroy the first order archetype.

Vertical disruption, corresponding to the customary description of first order configural properties from literature (Bentin, Allison, Puce, Perez, & McCarthy, 1996; Johnson, Dziurawiec, Ellis, & Morton, 1991), was as expected relevant as well. Finally, the deviation of single parts appears to be relevant only with respect to an axis defined by their position (eyes on *l_e_*, mouth and *l_nm_*) or somewhat related to it (nose and *l_pe_*, which in the archetype are aligned and superimposed). This may be due to the fact that those variations are local, and as such it would be more difficult to perceive alterations related to axes that are distant from the single parts.

Our results confirm that first order relational configural properties play an important role in face detection. Our mathematical formalization of faceness can be used to construct stimulus sets exhibiting varying degrees of faceness on a continuous scale, thus opening possibilities for further investigation of the face processing network. Until now faceness had mostly been modulated categorically, with a simple binary class face/non-face. The only option to have a continuously varying faceness score has been manipulating lower level properties to make the face less/more visible. Examples of this are filtering the image or scrambling its 2D Fourier transform phase. The usefulness of this kind of process towards the understanding of face detection is limited, considering that there is no control on how stimuli vary with respect to the space of high-level features extracted by the brain. This problem is especially evident when attempting to measure the degradation of face-like images using those methods. Typically, the absolute value of degradation for a face-like image is computed measuring the distance of it from a face template. In case of an average image, this is a mean squared error of the pixel values; for specific configurations it is instead the sum of the Euclidean distances of each part from the archetypal positions. Both of these measures fail to replicate important invariance properties of the face detection process: they are not invariant to size, featural properties, second order configural properties, position in the field of view and inversion.

Our approach, on the other hand, retains all these invariance properties (see supplementary figure 1) and allows for continuous modulation of those high-level properties relevant for face detection; that is, first order configural properties. Our model thus offers the opportunity to address hitherto unanswered questions regarding face detection and configural processing that were neglected due to a lack of rigorous mathematical formalizations (Maurer et al., 2002). Specifically, while the relevance of FOC properties for face detection has been shown in multiple studies, using event-related potentials (ERP) and functional magnetic resonance imaging (fMRI) (Ghuman et al., 2014; Maurer et al., 2002), the lack of a formalization of the properties that describe the FOC (and hence also the faceness) of a stimulus limited the ability to pinpoint the neural mechanism involved in face detection. Studies employing ERP and fMRI have suggested the fusiform face area (Ghuman et al., 2014) as well as the amygdala and superior temporal sulcus to be involved in face detection (Golarai et al., 2015). However, the unique contribution of each of these regions to face detection as well as their interactions remains an open question which requires the kind of continuous control over stimuli proposed here.

Nevertheless, the model we present here comes with some intrinsic limitations of its own. First of all, the use of limited featural information in the representation of the face parts, reduced to simple rectangles, necessitates empirically defined limits of deviation for the configurations to ensure that rectangles designated as a specific face part are indeed recognized as such. A more complete description would define all the features we formalized for all possible combinations of associations between face parts and rectangles. This would, however, multiply the number of features to consider and hence the dimensionality of the problem. Another limit lies in the fact that the distance measures are relative to the size of the canvas in which the face parts are disposed rather than to the size of the face itself. This problem is not trivially solvable because it is difficult to define a measure of the size of the face when its boundary is absent or unknown. Thanks to the reduction of the feature space based on our empirical findings, these issues might be more readily addressed by future studies to further improve our formalization.

In conclusion, our experimental findings confirmed the validity of our mathematical formalization of first order configural properties related to faceness and allowed us to identify a small set of maximally informative features for face detection, which we hope will aid future studies in localizing and identifying the neural mechanisms underlying face detection.

## 6. Acknowledgements

We also wish to thank Anna Sagana for her comments on this manuscript. This project was funded by the European Union’s Horizon 2020 research and innovation programme under the Marie Sklodowska-Curie grant agreement No 641805; MS and RG were also funded by the European FET Flagship project ‘Human Brain Project’ FP7-ICT-2013-FET-F/604102. Declarations of interest: none.

---

**Supplementary figure 1:**
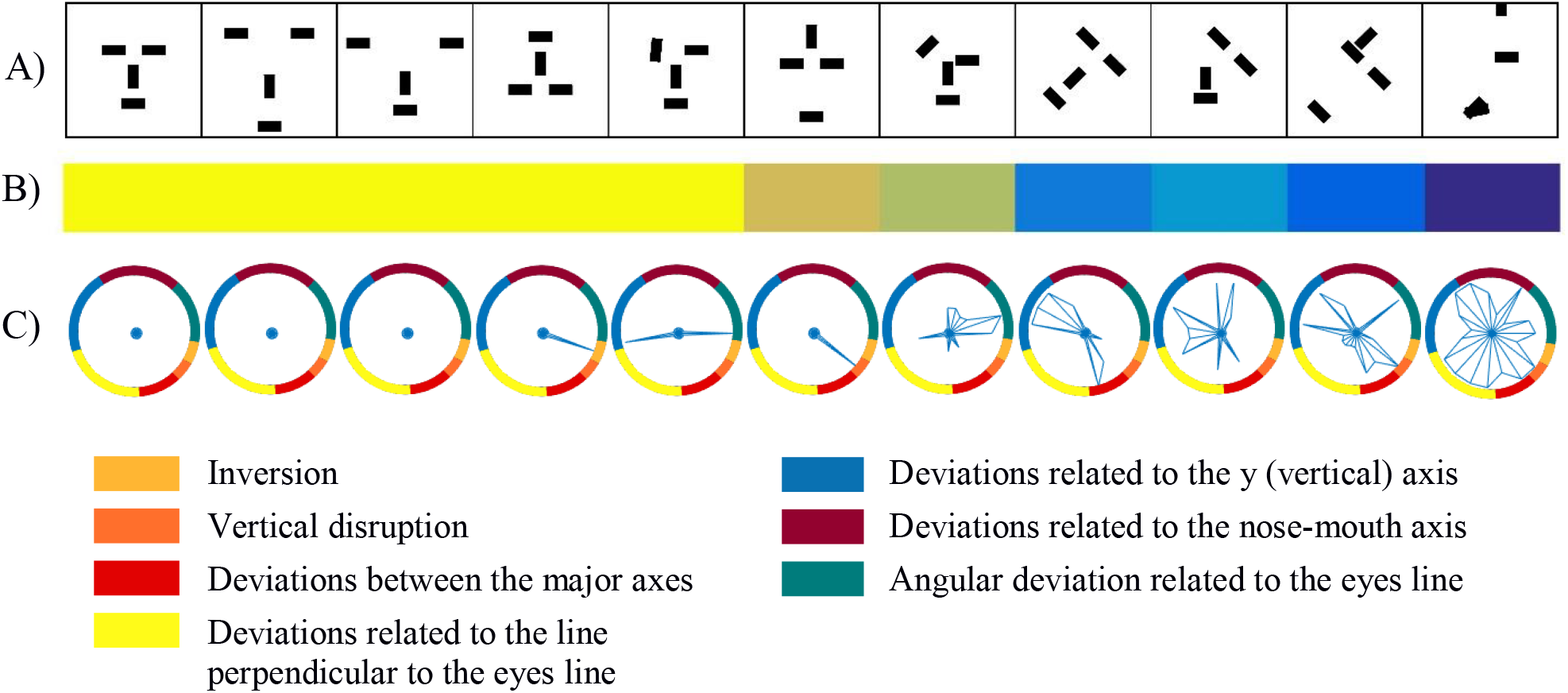
Exemplary model properties. (A) A range of configurations. (B) Glyph graphs of the feature values (the further the line extends from the center, the larger the feature value) in which features are divided by color. (C) *Faceness* value computed using the model determined using the *β*_2_ based stepwise regression.

## Appendix A: Weighted sampling method

Stimulus choice in the Likert experiment is constrained by several requirements:

1. Face parts must be identifiable
2. The distribution of feature values must be multivariate normal
3. Features must be independent (i.e. the covariance matrix must be diagonal)
4. Faceness (in terms of low global deviation from the archetype) must be normally distributed

Considering that the feature space is19-dimensional computed over the 12-dimensional domain in which the configural matrices are defined and that face parts are not recognizable, it is not feasible to use an optimization algorithm for finding a set of configurations fulfilling all criteria. Instead, we opt for generating a large set of configurations from which we can sample. The sampling procedure is weighted such that individual feature as well as faceness distributions across the selected configurations match pre-defined probability mass functions.

In what follows we describe how the initial set of configurations is generated and how the sampling weights are determined. To generate the initial set of configurations from which we sample the eventual stimulus set, we start with the configural matrix describing a single archetypal configuration (feature_i_(*m*) = 0, ∀i). Based on this we define, for each part, a set of positions along a range of dislocations which are sufficiently small for the part to be identifiable. With these restrictions in place, we compute the matrices determined by all possible combinations of those values, leading to a set of ~10^9^ configural matrices **m**.

This set enables us to estimate the multivariate probability mass function over the feature space. To that end, we discretize each feature’s values into 10 equidistant bins and compute histograms for all features (with the exception of vertical disruption which only takes on two values). More formally, we assign to each configuration matrix **m** an array ***b*** of 19 elements which indicates for each feature the bin into which it falls:

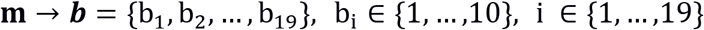

The array ***b*** thus realizes a discrete multivariate random variable B. In total, B has 19 elements B_i_, discrete random variables with values in {1, …,10} indicating the bin for each feature separately.

The probability mass function of each element B_i_ can be described as the ratio between the number of configurations for which the feature fell into a bin, *n_b_i__*, and the total number of configurations, *n_c_*:

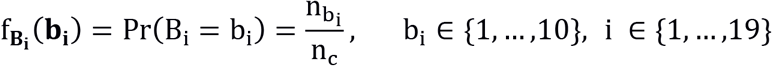

If we assume that variables B_i_ are independent, the conjunct probability mass function is given by the product of the probability mass functions of the individual elements:

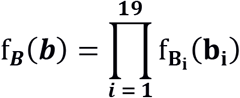

This means that the probability that a configuration can be described by an array ***b***, i.e. the same distribution across bins for all features, is given by the product of the probabilities of observing that distribution over bins in each feature:

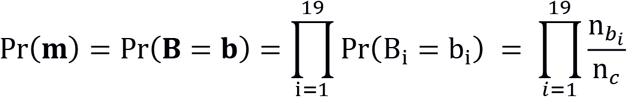

Given this, it is possible to sample a subset of **m** whose distribution over bins is uniform. This can straightforwardly be achieved using the reciprocal of Pr(**m**) as weight for the sampling:

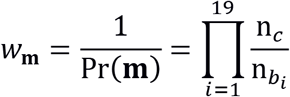

While a uniform distribution might seem like a good choice, ensuring an equiprobable distribution of feature values, it leads to sampling many configurations with extreme levels of distortion. For this reason, we prefer a Gaussian distribution to control the spread of values around pre-defined means. Specifically, we want individual features to be distributed according to N(μ = 0, σ = 2.5) and the normalized deviation of global faceness to be distributed as *N*(*μ* = 1.5, *σ* = 1). Considering that, by definition, the feature values and faceness can only be positive, both distributions are considered only in their positive part. To ensure that our stimulus set will follow the desired Gaussian distributions, we sample 10 equidistant values from reference Gaussian distributions, in the domain {0,5}, and multiply those values by the previously established weight of the corresponding bin index

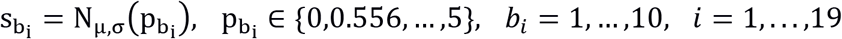

The weight for a specific configuration matrix then becomes:

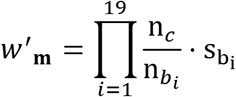

Finally, because the features are not truly independent and because we want to control the size of the stimulus set, we choose a subset of features of size *n* based on which, we compute sampling weights. In the case of our experiment, *n* = 9. As described previously, sampling is also weighted by global faceness. We obtain the corresponding weight by considering global faceness as another feature and apply the same principles described above.

## References

Bayliss, A. P., & Tipper, S. P. (2005). Gaze and arrow cueing of attention reveals individual differences along the autism spectrum as a function of target context. British Journal of Psychology, 96(1), 95–114. http://doi.org/10.1348/000712604X15626

Bentin, S., Allison, T., Puce, A., Perez, E., & McCarthy, G. (1996). Electrophysiological Studies of Face Perception in Humans. Journal of Cognitive Neuroscience, 8(6), 551–565. http://doi.org/10.1162/jocn.1996.8.6.551

Brainard, D. H. (1997). The Psychophysics Toolbox. Spatial Vision, 10(4), 433–6. http://doi.org/10.1163/156856897X00357

Chang, L., & Tsao, D. Y. (2017). The Code for Facial Identity in the Primate Brain. Cell, 169(6), 1013–1028. http://doi.org/10.1016/j.cell.2017.05.011

Chen, Y., McBain, R., & Norton, D. (2015). Specific vulnerability of face perception to noise: a similar effect in schizophrenia patients and healthy individuals. Psychiatry Research, 225(3), 619–24. http://doi.org/10.1016/j.psychres.2014.11.035

de Haas, B., Schwarzkopf, D. S., Alvarez, I., Lawson, R. P., Henriksson, L., Kriegeskorte, N., & Rees, G. (2016). Perception and Processing of Faces in the Human Brain Is Tuned to Typical Feature Locations. The Journal of Neuroscience, 36(36), 9289–9302. http://doi.org/10.1523/JNEUROSCI.4131-14.2016

Duchaine, B., & Yovel, G. (2015). A Revised Neural Framework for Face Processing. Annual Review of Vision Science, 1(1), 393–416. http://doi.org/10.1146/annurev-vision-082114-035518

Freiwald, W. A., Tsao, D. Y., & Livingstone, M. S. (2009). A face feature space in the macaque temporal lobe. Nature Neuroscience, 12(9), 1187–96. http://doi.org/10.1038/nn.2363

Ghuman, A. S., Brunet, N. M., Li, Y., Konecky, R. O., Pyles, J. A., Walls, S. A., … Richardson, R. M. (2014). Dynamic encoding of face information in the human fusiform gyrus. Nature Communications, 5, 1–10. http://doi.org/10.1038/ncomms6672

Gilad-Gutnick, S., Yovel, G., & Sinha, P. (2012). Recognizing Degraded Faces: The Contribution of Configural and Featural Cues. Perception, 41(12), 1497–1511. http://doi.org/10.1068/p7064

Golarai, G., Ghahremani, D. G., Eberhardt, J. L., & Gabrieli, J. D. E. (2015). Distinct representations of configural and part information across multiple face-selective regions of the human brain. Frontiers in Psychology, 6(NOV), 1–13. http://doi.org/10.3389/fpsyg.2015.01710

Golarai, G., Grill-Spector, K., & Reiss, A. L. (2006). Autism and the development of face processing. Clinical Neuroscience Research, 6(3), 145–160. http://doi.org/10.1016/j.cnr.2006.08.001

Hadjikhani, N., Kveraga, K., Naik, P., & Ahlfors, S. P. (2009). Early (M170) activation of face-specific cortex by face-like objects. Neuroreport, 20(4), 403–7. http://doi.org/10.1097/WNR.0b013e328325a8e1

Jiang, F., Dricot, L., Weber, J., Righi, G., Tarr, M. J., Goebel, R., & Rossion, B. (2011). Face categorization in visual scenes may start in a higher order area of the right fusiform gyrus: evidence from dynamic visual stimulation in neuroimaging. Journal of Neurophysiology, 106(5).

Johnson, M. H., Dziurawiec, S., Ellis, H., & Morton, J. (1991). Newborns’ preferential tracking of face-like stimuli and its subsequent decline. Cognition, 40(1), 1–19. http://doi.org/10.1016/0010-0277(91)90045-6

Kanwisher, N., & Barton, J. J. S. (2011). The Functional Architecture of the Face System: integrating Evidence from fMRI and Patient Studies. Oxford University Press. http://doi.org/10.1093/oxfordhb/9780199559053.013.0007

Kleiner, M., Brainard, D., Pelli, D., Ingling, A., Murray, R., & Broussard, C. (2007). “What’s new in Psychtoolbox-3?”. Perception ECVP Abstract Supplement, 36(14), 1–16. Retrieved from https://nyuscholars.nyu.edu/en/publications/whats-new-in-psychtoolbox-3

Kobatake, E., & Tanaka, K. (1994). Neuronal selectivities to complex object features in the ventral visual pathway of the macaque cerebral cortex. Journal of Neurophysiology, 71(3), 856–867. http://doi.org/10.1038/476265a

Liu, J., Harris, A., & Kanwisher, N. (2002). Stages of processing in face perception: an MEG study. Nature Neuroscience, 5(9), 910–916. http://doi.org/10.1038/nn909

Macchi Cassia, V., Kuefner, D., Westerlund, A., & Nelson, C. A. (2006). A behavioural and ERP investigation of 3-month-olds’ face preferences. Neuropsychologia, 44(11), 2113–2125. http://doi.org/10.1016/j.neuropsychologia.2005.11.014

Maurer, D., Grand, R. Le, & Mondloch, C. J. (2002). The many faces of configural processing. Trends in Cognitive Sciences, 6(6), 255–260. http://doi.org/10.1016/S1364-6613(02)01903-4

Mondloch, C. J., Le Grand, R., & Maurer, D. (2002). Configurai face processing develops more slowly than featural face processing. Perception, 31(5), 553–566. http://doi.org/10.1068/p3339

Pascalis, O., & Kelly, D. J. (2009). The origins of face processing in humans phylogeny and ontogeny. Perspectives on Psychological Science, 4(2), 200–209. http://doi.org/10.1111/j.1745-6924.2009.01119.x

Pelli, D. G. (1997). The VideoToolbox software for visual psychophysics: transforming numbers into movies. Spatial Vision, 10(4), 437–42. Retrieved from http://www.ncbi.nlm.nih.gov/pubmed/9176953

Sheehan, M. J., & Nachman, M. W. (2014). Morphological and population genomic evidence that human faces have evolved to signal individual identity. http://doi.org/10.1038/ncomms5800

Sun, H.-M., & Balas, B. (2014). Face features and face configurations both contribute to visual crowding. Attention, Perception & Psychophysics, 77, 508–519. http://doi.org/10.3758/s13414-014-0786-0

Tanaka, J. W., & Gordon, I. (2011). Features, Configuration, and Holistic Face Processing. Oxford University Press. http://doi.org/10.1093/oxfordhb/9780199559053.013.0010

Tanaka, J. W., & Sengco, J. a. (1997). Features and their configuration in face recognition. Memory & Cognition, 25(5), 583–592. http://doi.org/10.3758/BF03211301

Tanaka, K. (2003). Columns for complex visual object features in the inferotemporal cortex: clustering of cells with similar but slightly different stimulus selectivities. Cerebral Cortex (New York, N.Y.: 1991), 13(1), 90–99. http://doi.org/10.1093/cercor/13.1.90

Tsao, D. Y., & Livingstone, M. S. (2008). Mechanisms of Face Perception. Annual Review of Neuroscience, 31(1), 411–437. http://doi.org/10.1146/annurev.neuro.30.051606.094238

